# Genomic diversity, population structure and accessory genome analysis of *Pasteurella multocida*: New Insights into host adaptation and disease specialization

**DOI:** 10.1101/2020.11.30.403022

**Authors:** Dennis Carhuaricra, Raquel Hurtado, Luis Luna, Raúl Rosadio, Jane C. Wheeler, Vasco Azevedo, Lenin Maturrano

## Abstract

*Pasteurella multocida* is a multi-host pathogen that infects a wide spectrum of domestic and wild animals including humans. Despite its impact on health and economics, *P. multocida* is considered an enigmatic pathogen and the genetic basis of its pathogenicity and host adaptation still remains unclear. Here we present a detailed genomic framework based on 336 whole-genome sequences of *P. multocida* isolates from different animal species and countries. Our data provide genomic support of the existence of two very divergent phylogroups (PmI and PmII), which present a barrier to homologous recombination suggesting genetic isolation. Additionally, a *torCAD* operon, which reduces TMAO (trimethylamine *N*-oxide) to produce energy during bacterial anaerobic respiration, is present only in PmI and can act as a hypothetical driver of niche segregation between phylogroups. The PmI phylogroup harbors strains that infect a wider range of hosts than PmII, and shows a highly diverse phylogeny and accessory genome. We identified nine clonal lineages for PmI, seven of which are associated with specific hosts or diseases and contain distinct accessory gene pools that can confer ecologically relevant phenotypes. We found differential presence of a trehalose metabolism operon in the bovine lineage associated with pneumonic pasteurellosis; while, citrate, L-arabinose, L-fucose, and D-allose operons are only present in avian lineages. These findings suggest that alternative metabolic pathways may facilitate the establishment of *P. multocida* during host colonization in the early stages of infection promoting the adaptation of *P. multocida* lineages to certain hosts.

## Introduction

*Pasteurella multocida* is considered a versatile bacterium capable of infecting a wide range of animal hosts producing different clinical manifestations in each one (Wilkie et al., 2012). The myriad of diseases produced by *P.multocida* in domestic and wild animals includes fowl cholera in poultry and wild birds, hemorrhagic septicemia in bovines (mainly in Asia), pneumonic disease (“snuffles”) in rabbits, atrophic rhinitis in pigs and pneumonia in cattle, pigs, alpacas and sheep (Rosadio et al., 2011; Shivachandra et al., 2011; Wilkie et al., 2012). *P. multocida* can also produce uncommon infections in humans through animal bites or scratches mainly from cats and dogs (Wilson and Ho, 2013). Despite the impact on health, economics and conservation, *P. multocida* has been considered an “enigmatic pathogen” due to poor understanding of the genetic factors underlying its varied pathogenicity (Boyce et al., 2012; Wilkie et al., 2012).

Five capsular types (A, B, D, E and F) of *P. multocida* have been classified by serotyping, as well as 16 somatic serotypes based on lipopolysaccharide (LPS) antigens (Townsend et al., 2001; Harper et al., 2012) and recently have been reclassified into 8 LPS genotypes L1-L8 using multiplex-PCR (Harper et al., 2015). Both the capsular types and the LPS serotypes are considered important virulence factors; for example, capsular type B causes hemorrhagic septicemia in bovines (Boyce and Adler, 2000) and type A causes fowl cholera in poultry (Chung et al., 2001). Acapsular and mutant LPS strains are much less virulent and are completely attenuated (Harper et al., 2004). Other important virulence factors that contribute to *P. multocida* pathogenesis are adhesins, toxin A, iron metabolism related proteins and outer membrane proteins (Harper et al., 2006).

Epidemiological studies using multilocus sequence-typing (MLST) have revealed some clonal lineages associated with specific diseases (Hotchkiss et al., 2011a). For example, the clonal complex (CC) CC13 has been associated with bovine respiratory disease worldwide (Hotchkiss et al., 2011b), while CC122 is associated to hemorrhagic septicemia in wild and domestic bovines mainly in Asia (Petersen et al., 2014), suggesting the existence of lineages with a predilection to produce certain clinical manifestations in some hosts. The ST9 lineage is recognized as generalist and associated with rabbits, poultry and pigs (Hotchkiss et al., 2011a).

In recent years, massification and cheaper sequencing techniques have allowed the sequencing of many *P. multocida* whole genomes worldwide (Peng et al., 2019; Hurtado et al., 2020). These abundant data reveal high genetic diversity with the abundant presence of mobile genetic elements including phages, genomic islands, plasmids and/or ICE carrying antimicrobial resistance genes (Michael et al., 2012; Moustafa et al., 2015; Zhu et al., 2019). The availability of whole genome sequences has made it possible to carry out comparative genome studies that have revealed an open pangenome for *P. multocida* with an important impact of horizontal gene transfer during their evolution (Hurtado et al., 2018; Zhu et al., 2019). Phylogenetic analyses of the entire genome have reported the existence of lineages consistent with the MLST system and the LPS genotype, suggesting the presence of genetic structure in *P. multocida* populations (Peng et al., 2018; Zhu et al., 2019). Comparative analyses of a few genomes have described the presence of genes associated with certain hosts such as L-fucose and citrate operons in virulent isolates from poultry (Johnson et al., 2013); whereas trehalose metabolism related genes have been reported in cattle isolates (Hurtado et al., 2018). It is clear that broader data are needed to improve understanding of the high *P. multocida* diversity, evolution and host adaptation (Peng et al., 2019; Hurtado et al., 2020).

In the present work, we carry out a broad analysis of the genomic diversity, population structure and accessory genome of 336 *P. multocida* genomes, including 5 new genomes isolated from alpacas in Peru. This collection represents a wide range of hosts and geographic regions and provides high-resolution evidence of *P. multocida* adaptation to hosts and diseases.

## Methods

### *Pasteurella multocida* isolates, sequencing and publienome sequences

Five *Pasteurella multocida* strains isolated from alpacas with acute pneumonia in Peru were selected from the collections of the Molecular Biology and Genetics Laboratory (Faculty of Veterinary Medicine, Universidad Nacional Mayor de San Marcos) for sequencing. The strains were previously characterized by our group (Rímac et al., 2017). These isolates were grown in LB broth overnight at 37°C and genomic DNA was extracted using the PureLink Genomic DNA Mini kit (Invitrogen). Illumina sequencing was performed using Nextera XT DNA Library Preparation Kit and Miseq System yielding 251-bp paired-end reads. Read quality was checked with fastQC and trimmed using Trimmomatic (Bolger et al., 2014). *De novo* assembly was performed using SPAdes 3.14.1 (Bankevich et al., 2012). Contig sequences are deposited in GenBank under assembly accession numbers: JACDXE000000000-JACDXI000000000.

Public genome sequences of P. *multocida* were retrieved from the Genome database (ftp://ftp.ncbi.nih.gov/genomes/) and European Nucleotide Archive (ENA) FASTQ database (https://www.ebi.ac.uk/ena/browse/download) in September 2019. *de novo* assembly of the raw data was carried out using the same pipeline above. Our final genome dataset consists of 340 complete *P. multocida* genomes including the five new alpaca genome sequences (Supplementary Data Table S1).

### Genetic diversity and Pangenome analyses

Average Nucleotide Identity (ANI) analysis with python pipeline Pyani (http://widdowquinn.github.io/pyani/) was performed for the 340 genomes to confirm P. *multocida* species membership. Four highly divergent sequences (<94% identity) were eliminated and a total of 336 *P. multocida* genome sequences were annotated with Prokka (Seemann, 2014). The GFF3 annotation format assemblies were input to Roary (Page et al., 2015) to construct the pangenome with default settings of 95% identity and coverage. Pangenome accumulation curves were generated for P. *multocida* using vegan package in R from the presence/absence matrix of orthologous groups retrieved from Roary.

### Phylogroup and recombination analysis

The core genome alignment retrieved from Roary (570502 aligned position) was used to generate a NeighbourNet split network with the uncorrect P distance parameter with SplitsTree4 (Kloepper and Huson, 2008). Genetic divergence between PmI and PmII phylogroups was calculated in R. PCA analysis of the accessory genome of different phylogroups was performed with *ggfortify* package. Recombinant regions were identified running Gubbins with default parameters (Croucher et al., 2015). A network analysis of recombinant regions identified by Gubbins between phylogroups was performed using *igraph* package considering edge as 10% of recombination regions shared.

### Phylogenomics and population structure of PmI phylogroup

Core genome alignment from PmI phylogroup was filtered using Gubbins to remove high SNP density regions which indicate putative recombinant events using 10 iterations. To estimate a phylogenetic tree, we used RAxML (Stamatakis, 2014) with the GTR+Gamma model on core gene alignment stripped of recombination with 1000 bootstrap replicates. Phylogenetic trees were visualized using *ggtree* package on R.

Identification of the major population groups was determined using hierBAPS. Two levels with a maximum of 15 populations were set as the initial condition for rhierBAPS (Tonkin-Hill et al., 2018) to infer the population structure of *Pasteurella multocida* genomes using a maximum-likelihood approach. We also performed the multilocus sequence type (MLST) analysis for *P multocida* lineage genotyping with the *mlst* pipeline from T. Seemman (https://github.com/tseemann/mlst) using the RIRDC scheme database.

### Accessory genome analysis of *P. multocida* PmI phylogroup

We used the presence/absence matrix of accessory genes recovered from Roary to calculate a pairwise distance matrix using Jaccard function in R. Then, this matrix was transformed into a bi-directional network using igraph package with edges indicating at least 70% of accessory genes shared between isolates (nodes) and edge length weighted according to the proportion of genes shared (shorter edges means more accessory genes shared between genomes). We also performed a PCA analysis in the same matrix to assess clustering between isolates from different hosts using ggfortify package.

### Identification of gene clusters associated to specific host lineages

Cluster defining accessory genes were identified with k-pax2 (Pessia et al., 2015) by running Bayesian method on the accessory genome matrix with default parameters. Genes with high discrimination power for BAPS clustering were functionally annotated using the eggNOG database and the eggnog-mapper tool (Huerta-Cepas et al., 2017). Operons and differentially present regions were drawn using the online tool Gene Graphics (Harrison et al., 2018).

### Genotyping and distribution of virulence factors and antimicrobial resistance genes

In-silico genotypification for all *P. multocida* genomes was carried out identifying gene sequences used in PCR for LPS (L1-L8) and Capsular (A, B, D, E and F) typing (Townsend et al., 2001; Harper et al., 2015) in the genome sequences of the 336 isolates. The same strategy was used to identify the presence and integrity of known virulence genes present in *P. multocida*. Antimicrobial resistance genes present in the genome sequences were identified using *abricate* (https://github.com/tseemann/abricate) against the CARD and ARG-ANNOT databases (Gupta et al., 2014; Alcock et al., 2020).

## Results

### Global *Pasteurella multocida* genomes collection

To investigate the population structure and genomic evolution of *P. multocida* we analyzed 340 whole genome sequences including 5 new sequences isolated from alpacas. These sequences belong to 10 different hosts from 20 different countries across 5 continents (Supplementary Data Table S1). The Average Nucleotide Identity (ANI) was calculated for all possible pairs of genomes to evaluate the genetic diversity and species membership across all isolates considering 95% as species delimitation. This analysis revealed four genomes with an ANI < 95% compared to other genomes and these were removed from subsequent analysis (Supplementary Fig. 1A). The remaining 336 *P. multocida* genomes belong to a single species (ANI > 95%) and were classified in 6 groups corresponding to host (alpaca, bovine, avian, swine, rabbit and human) and a category “other” with no more than 2 isolates per animal (mice, dogs, monkeys and sheep) (Supplementary Fig. 1B).

### Pangenome and phylogenomics of *P. multocida* support two genetically divergent subpopulations

Pangenome size was 13550 genes for the 336 *P. multocida* genomes. The gene accumulation curve (Supplementary Fig. 2A) shows an open pangenome indicating that new genes will continue increasing with additional genome sequences. The core genome of *P. multocida* harbors a total of 776 genes, with a sequence length of 570502 Mpb and containing 44868 SNPs. We identified two phylogroups with 100% bootstrap support using split network and maximum likelihood phylogenetic analyses (Fig 1A, 1D) which we named PmI (282 genomes) and PmII (54 genomes). The highly structured phylogeny of *P. multocida* shows two phylogroups with a nucleotide divergence of 3-4% (calculated across the genome) (Fig 1B), suggesting the absence of gene flow between these phylogroups. To confirm this, we explore recombination patterns across the genome which revealed a strong barrier to homologous recombination between PmI and PmII (Fig 1C).

**Figure 1.**
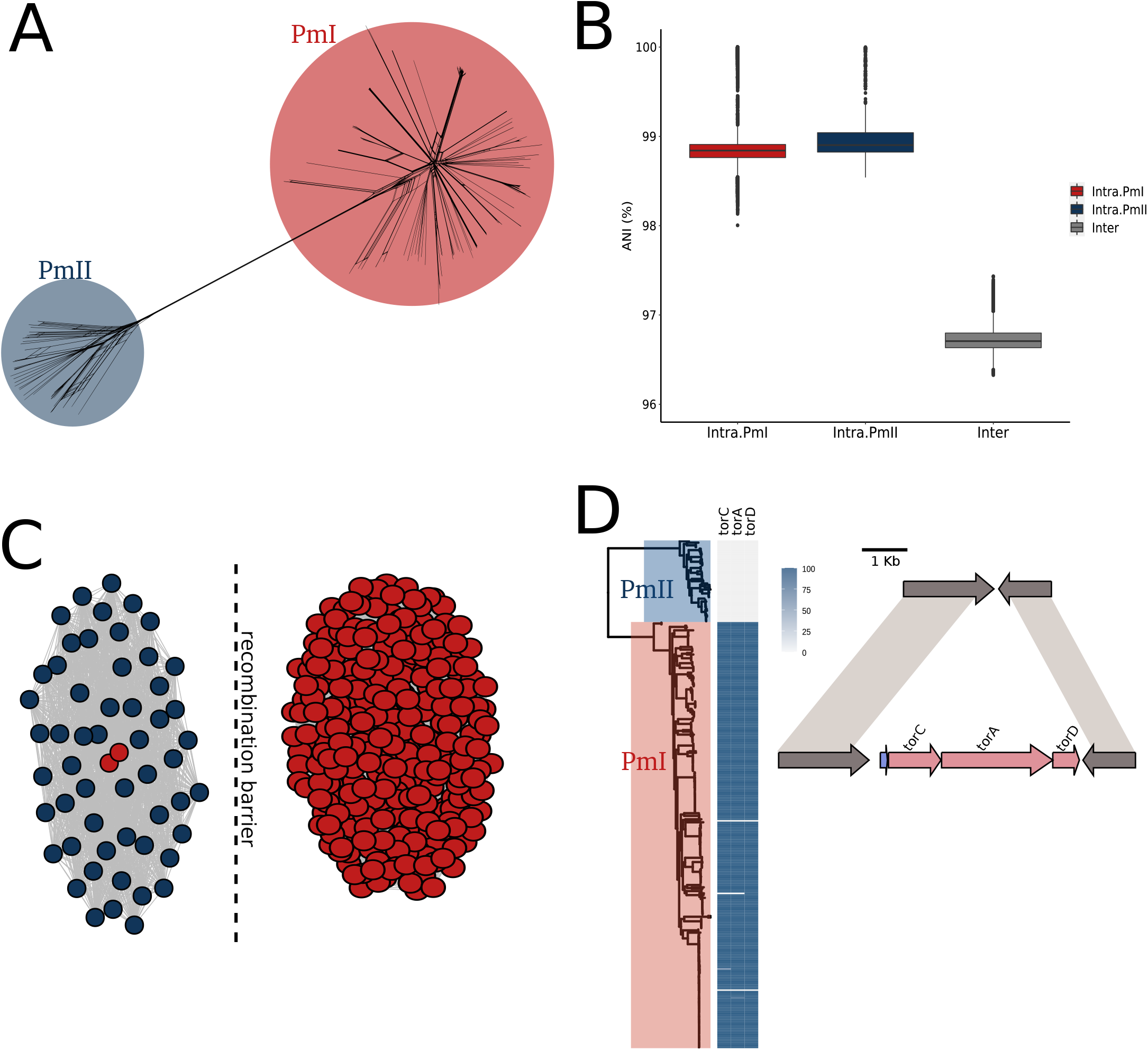
Phylogeny and recombination of ecologically distinct *P. multocida* PmI and PmII phylogroups. (A) Split network of 336 *P. multocida* genomes with phylogroups highlighted. (B) Boxplots showing ANI values calculated intra and inter phylogroups. Inter-phylogroups ANI distance is around 96-97% (C) Network analysis of shared recombination blocks (edges) between PmI and PmII phylogroups. each edge indicates at least 10% of recombination blocks shared between isolates. A clear barrier to recombination is drawing between phylogroups (D) Maximum likelihood tree of 336 *P. multocida* genomes (left) coupled to heatmap of presence/absence for *torC, torA and torD* genes showing that torCAD operon is present only in isolated from PmI phylogroup.

The analysis of accessory gene abundance among these phylogroups reveals more genes in PmI than PmII (Supplementary Fig. 2B). PmII shows an apparent host restriction with 47 of 54 sequences from avian and 6 from human host samples from the USA and Europe. In contrast, PmI harbor strains from all host species in this study including bovine, avian, swine, alpaca and human; a wide range of hosts demands more gene repertoire. Additionally, principal component analysis (PCA) of accessory gene content clearly distinguished the two phylogroups (Supplementary Fig. 2C). Together, this data provides whole-genome support for two separate lineages of *P. multocida* with considerable genetic isolation constituting discrete bacterial populations that are evolving independently.

Our data indicate that although PmI and PmII phylogroups possess the capacity to infect avian hosts, no evidence exists of differential metabolism, pathogenicity or barrier for recombination between isolates from these phylogroups. But differential gene content between these lineages can be important to niche specialization. Manual inspection of gene content revealed that the operon *torCAD* is present in PmI and absent in isolates from PmII (Fig 1D). The *torCAD* operon reduces TMAO (trimethylamine *N*-oxide) to TMA (trimethylamine) to produce energy in *Escherichia coli*. In bacterial cells TMAO acts as an electron acceptor during anaerobic respiration (Méjean et al., 1994). The *torCAD* operon could be responsible for differential physiological capacities between these phylogroups and contribute to niche specialization.

### Population structure and phylogenomics of PmI phylogroup show lineages strongly associated to host and disease

We constructed a maximum likelihood phylogeny of 282 genomes from PmI phylogroup based on a single copy gene of the core genome alignment. We determined 35660 single-nucleotide variable sites from the alignment and removed regions with elevated densities of base substitutions which indicate putative recombinant events with gubbins. The PmI phylogeny revealed a star-like population structure with many lineages strongly supported by bootstrap and some lineages showing predilection to a specific host and geographical distribution (Fig 2A).

**Figure 2.**
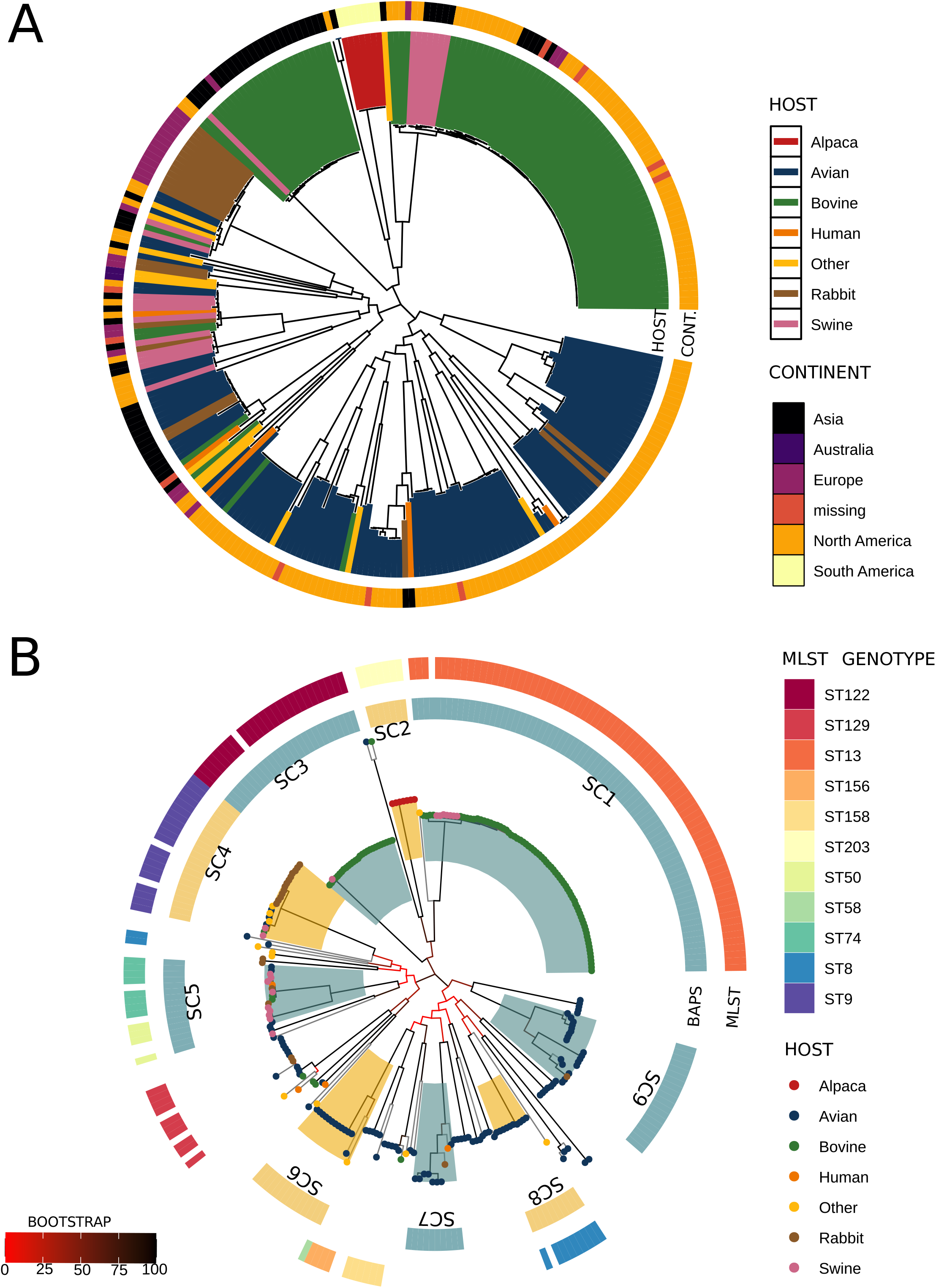
Phylogenomics and population structure of the *P. multocida* PmI phylogroup. (A) core genome phylogenomics of 282 PmI phylogroup genomes constructed using the maximum likelihood method showing host and geographic origin. Some clades are mainly associated to specific host. (B) ML tree and population structure of PmI phylogroups shows nine monophyletic sequence clusters (SCs) associated to MLST-clonal complex genotypes and host and disease predilection for seven SC (Table 1). Branch colors indicate bootstrap support according to the legend provided in the figure. hierBAPS sequence clusters are highlighted in alternating colors and labeled.

The population structure analysis revealed by hierBAPS identified 11 primary sequence clusters (SC). Of these, SC1 to SC9 were monophyletic clades, while the remaining SC10 and SC11 contains sequences not monophyletically distributed (Fig 2B). The monophyletic SCs were consistent with phylogenetically defined lineages with strong bootstrap support and most SCs contained isolates with specific MLST genotypes (Clonal Complex) (Fig 2B). SC1 (CC13) is composed mainly of bovine isolates of serotype A from different continents, while SC2 (CC203) is associated with alpaca isolates from South America; SC3 (CC122) is associated with bovine isolates of serogroup B that is responsible for hemorrhagic septicemia in Asia. SC4 (CC9) and SC5 (CC50 y CC74) are multi-host lineages. SC6, SC7, SC8 (CC8) and SC9 are mainly associated with avian isolates (Table 1). These data show some lineages, such as SC1, have a predilection for a specific host and are distributed worldwide. Some lineages, such as SC2 and SC3, are associated with a specific hosts or diseases but this could be influenced by local isolation. In other cases, multiple lineages can be restricted to a unique host, or be able to infect multiple hosts. The phylogeny reveals a high diversification of *P. multocida* with many lineages showing a predilection to different hosts.

**Table 1:** Sequence clusters identified by hierBAPS with respective MLST, LPS and capsular genotypification.

| SEQUENCE CLUSTER | HOST PREDILECTION | ASSOCIATED DISEASE | MLST | LPS | CAPSULE |
| --- | --- | --- | --- | --- | --- |
| SC1 | Bovine and pigs | Pneumonia | CC13 | L3 | A |
| SC2 | Alpaca | Pneumonia | CC203 | L6 | A |
| SC3 | Bovine | HS | CC122 | L2 | B |
| SC4 | Multi-host | Snuffles, FA, pneumonia | CC9 | L3 | A and F |
| SC5 | Multi-Host | FC, AR, pneumonia | CC74 and CC50 | L6 | A and D |
| SC6 | Poultry | FC | - | - | A |
| SC7 | Mainly poultry | FC | - | L5-L3 | A and F |
| SC8 | Poultry | FC | CC8 | L3 | A |
| SC9 | Poultry | FC | - | L3 | A and F |

Some lineages were concordant with Capsular: LPS: MLST genotypes described by Peng et al. 2018. For example, SC1 lineage is characterized by A: L3: CC13 genotype; SC2 by A: L3: CC203 genotype; SC3 by B: L2: CC122 genotype; whereas SC4 by A/F: L3:CC9 genotype and SC5 by A/D: L6: CC74/CC50 genotypes (Table 1 and Supplementary Fig. S3).

### The Accessory genome shows coherence with core genome phylogeny and clonal lineage distribution

The analysis of 336 genomes by Roary yielded a set of 5152 genes representative of the accessory genome (genes present in 2%-98% of isolates). Supplementary Fig. S4 shows differential gene content across different phylogenetic lineages, in this context, we find a high correlation between the pangenome tree constructed from presence/absence matrix distance with the core genome tree (p = 0.001 by Mantel test). Together, these are evidence of a coherent core genome and accessory genome trajectories. To assess the accessory genome distribution in different isolates according to host, we performed Network and PCA analyses from the presence/absence matrix of accessory genes (Fig 3). Network analysis supports clonal lineage structure revealed by population structure analysis with core-genome clonal lineages sharing clusters of accessory genes. Host associated lineages SC1, SC2, SC3 are present in separate clusters, the same for generalist lineages SC4 and SC5. While avian lineages are connected in one network, SC6 is a separate cluster. This strongly points to the existence of a host-specific gene pool required for *P. multocida* host adaptation. This suggests that some accessory gene combinations may confer a specific host tropism with the capacity to infect a specific host.

**Figure 3.**
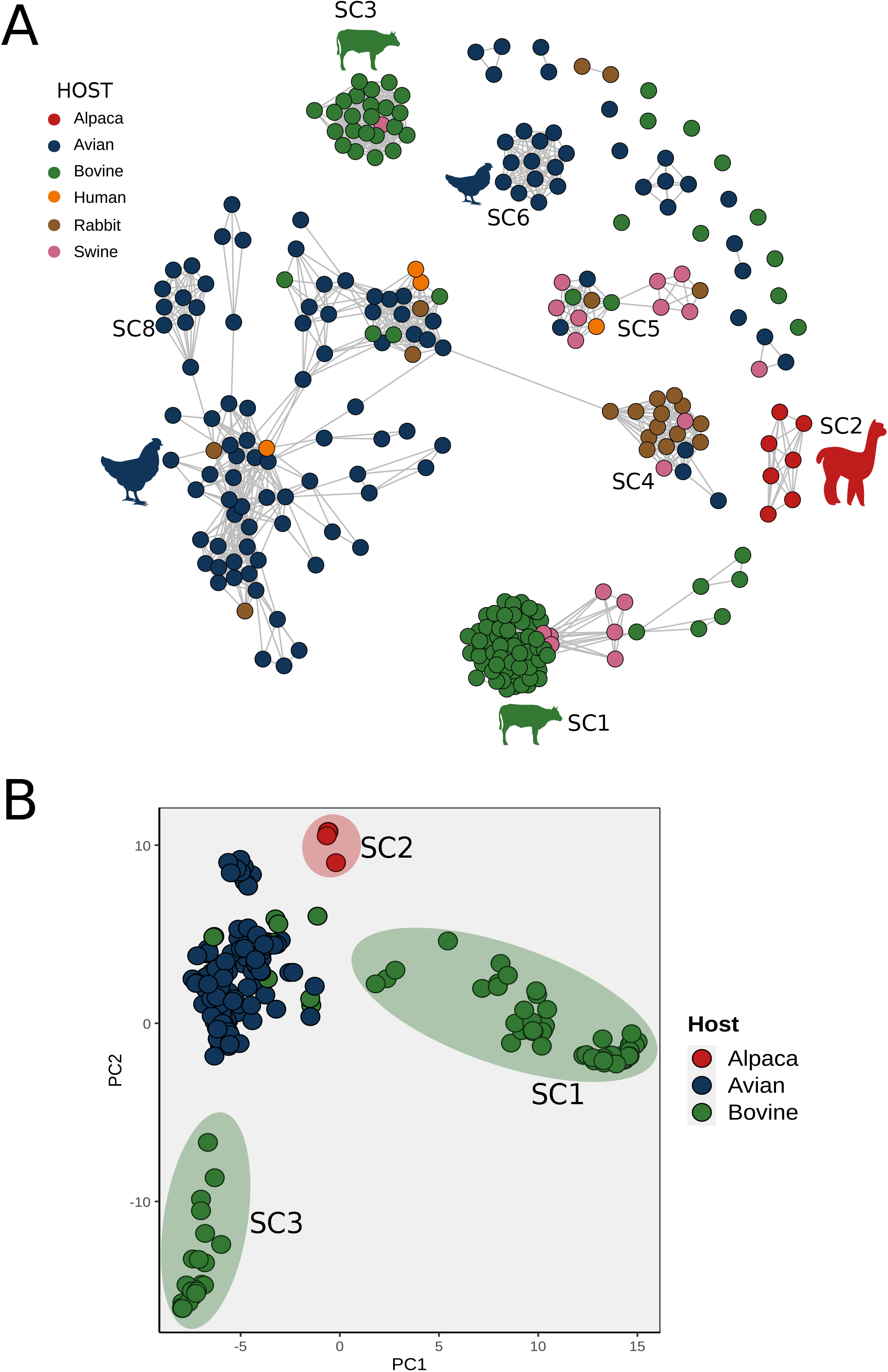
Network and PCA analyses of the PmI phylogroup accessory genome indicative clustering according to SC lineages and host and disease groups. (A) Network graph of 282 PmI isolates by correlation of 5152 accessory genes. Each node represents an isolate and is colored according host origin. The edge indicates > 70% shared genome content, with the length of the edges weighted by proportion of genes shared. The edges with < 70 % of accessory genes shared were removed. (B) PCA analysis of accessory genome between avian, bovine and alpaca isolates supporting the network graph findings above. Each point represents an isolate and also is colored according host-species origin.

### Differential gene pools are associate to specific host and disease lineages

We use a Bayesian probabilistic framework to detect those accessory genes with high discriminatory power to cluster isolates according to clonal lineage observed in network analysis. We identified 595 discriminatory genes principally from phage sequences and genes related to carbohydrate metabolism (Fig 4 and Table 2). We annotate all genes using eggNOG database independently for each clonal lineage. The SC1 lineage comprises serotype A isolates, mainly from bovine hosts with respiratory pneumonia, and harbors differentially an operon of trehalose (*treRBC*) absent in other lineages (Fig 5A). This *treRBC* operon can allow strains that carry it to import and metabolize this carbohydrate. A recent study has shown that a *Clostridium difficile* lineage has greater virulence than other lineages due to their ability to metabolize low concentrations of trehalose (Collins et al., 2018). Interestingly, the *ttrRSBCA* operon is absent in all strains of the SC1 lineage, but is present in all other lineages (Supplementary Fig. 5). The *ttrRSBCA* operon is known to play a role in anaerobic respiration in Salmonella during its colonization of the intestine (Fàbrega and Vila, 2013). Genes expressed by this operon use tetrathionate, which is normally produced in the cecal mucosa, as an electron acceptor. This advantageous ability is particularly important under the anaerobic growth conditions encountered in the intestinal mucus layer since it confers the opportunity to grow over fermenting commensal competitors (Winter et al., 2010).

**Figure 4.**
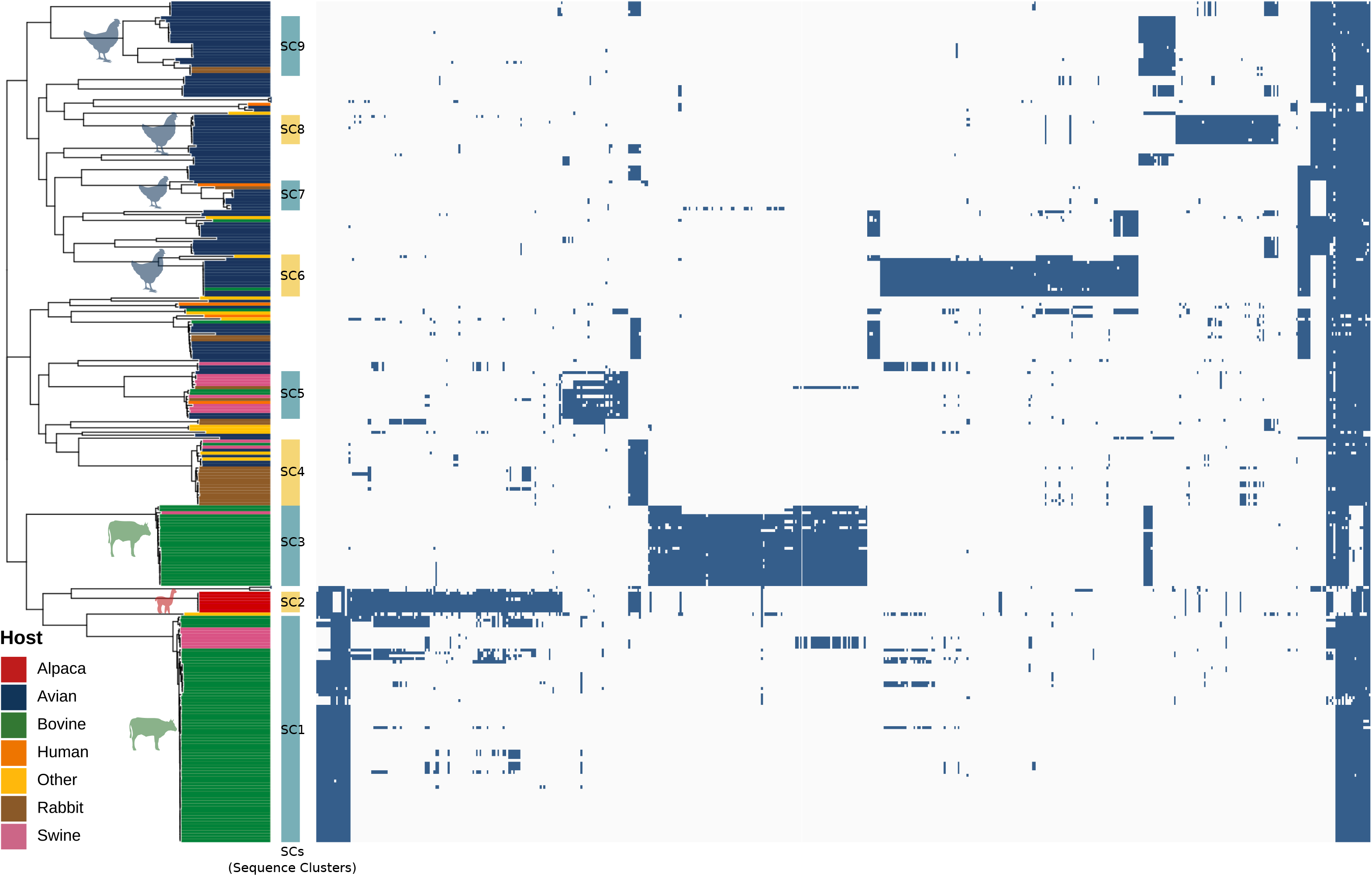
Heatmap of presence/absence discriminatory accessory genes identified by kpax2 Bayesian framework. Maximum likelihood tree of 282 *P. multocida* isolates from PmI phylogroup (left) with line colors representing host-species origins, and heatmap indicating presence (blue) or absence (white) of 595 accessory genes identified by Kpax2 as genes with more power to discriminate lineages clusters. Gene annotation of these genes revealed phage sequences and carbohydrate metabolism and transport genes (Table 2).

**Table 2:** List of accessory genes differentially present or absent in each lineage of *P. multocida*.

| SEQUENCE CLUSTER | HOST PREDILECTION | ASSOCIATED DISEASE | DIFFERENTIAL CLUSTER PRESENT | DIFFERENTIAL CLUSTER ABSENT |
| --- | --- | --- | --- | --- |
| SC1 | Bovine and pigs | Pneumonia | *treRBC* operon (trehalose metabolism) | *ttrRSBCA* operon (tretathionate metabolism) |
| SC2 | Alpaca | Pneumonia | 2 complete phage sequences,<br>1 plasmid sequence<br>1 phage with *toxA* gene | *hpaA*, *hpaBC* y *hpaG1G2EDF* (hydroxyphenylacetic metabolism) |
| SC3 | Bovine | HS | Capsule B locus<br>LPS locus<br>1 complete phage sequence<br>1 GI complete sequence | Capsule A locus |
| SC4 | Multi-host | Snuffles, FA, pneumonia | - | - |
| SC5 | Multi-Host | FC, AR, pneumonia | - | - |
| SC6 | Poultry | FC | *gatYZABCR* operon (galactitol metabolism)<br>cluster of 7 glucosyltransferase genes related-LPS metabolism<br>galactose operon<br>*alsRBACE* operon (D-Allose metabolism) | - |
| SC8 | Poultry | FC | *citCDEFG* operon (citrate metabolism)<br>CRISPR cluster genes<br>*tatD* gene<br>*FucPAUKIR* operon (L-Fucose metabolism) | - |
| SC9 | Poultry | FC | *FucPAUKIR* operon (L-Fucose metabolism) | - |

**Figure 5.**
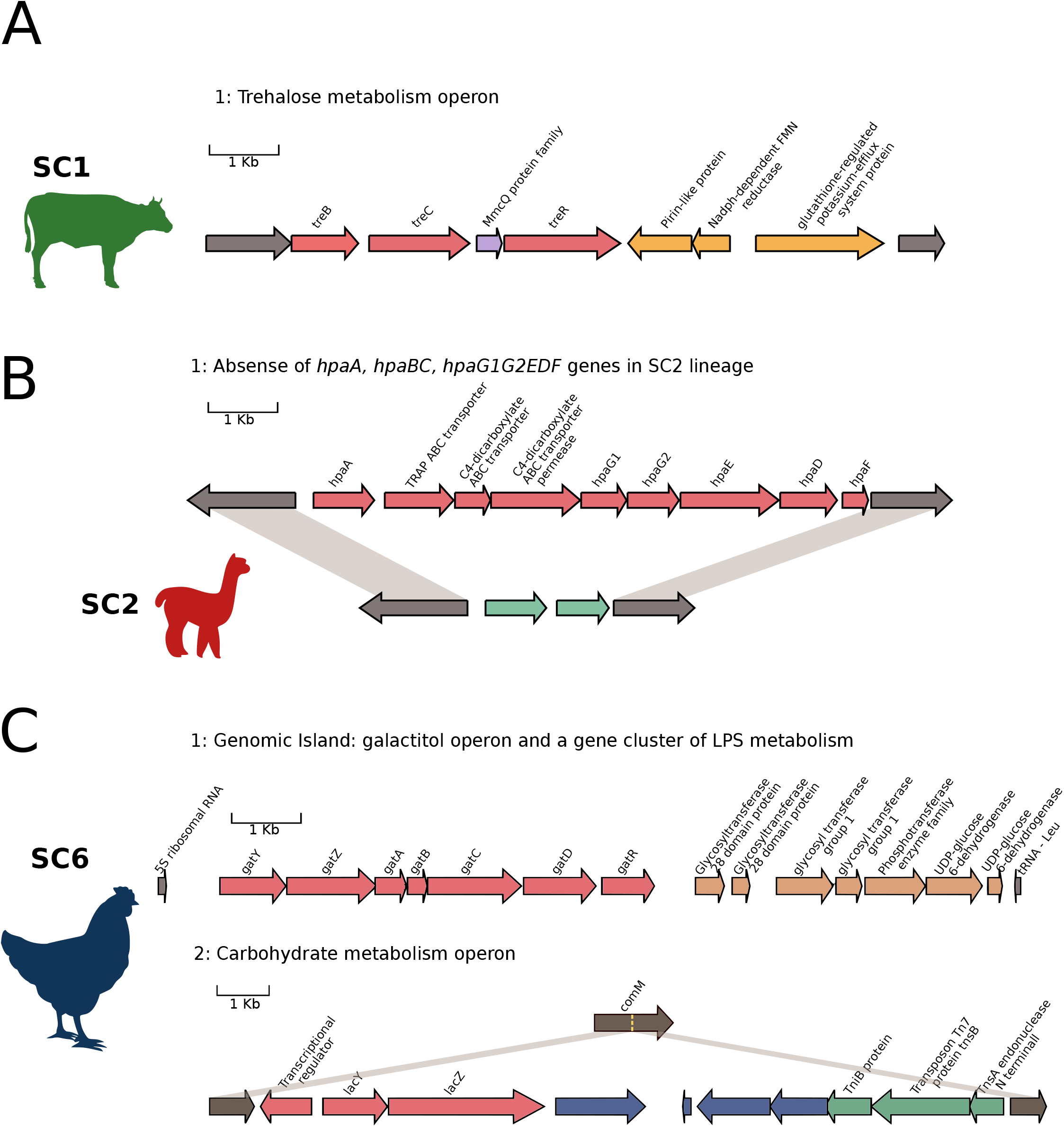
Gene annotation of accessory genes correlated to SC lineages and potentially implicated in host and disease adaptation. (A) gene annotation for a trehalose operon presents in SC1 lineage pneumonia associated). (B) In the SC2 lineage alpaca-pnumonia associated is absent a group of genes *hpaA, hpaBC y hpaG1G2EDF* related to hydroxyphenylacetic metabolism that are present in the other lineages. (C) Two gene clusters are presents in SC6 lineage poultry-Fowl cholera associated. First, a galatitol operon (*gatYZABCR*) with 7 genes related to glucosyltransferase function, maybe implicated in a new LPS from and second an operon with hyportetical galactose function inserted in the *comM* gene with transposable element genes.

The SC2 lineage, composed of alpaca isolates, contains 2 phage sequences, one with high similarity to the CIRMBP-0884 plasmid isolate from rabbits and a partial phage sequence containing the *toxA* gene (Table 2). As all 7 alpaca isolates are in contigs, it is not possible to know the complete sequence of mobile gene elements. Interestingly, alpaca isolates don’t have the cluster of 9 genes (operon *hpaA*, *hpaBC*, *hpaG1G2EDF*) related to utilization of hydroxyphenylacetic acid (HPA), all these genes are only absent in SC2 isolates (Fig 5B). Aromatic compounds are widely abundant in soil, water and animal gut and represent a common carbon source for many bacteria including commensal enteric as *E. coli* (Díaz et al., 2001). A study of porcine extraintestinal pathogenic *E. coli* (ExPEC) found that the virulent strain harbors this operon compared to commensal strains (Liu et al., 2015).

The hemorrhagic septicemia (HS) associated strains of SC3 lineage contain differentially the capsule B locus composed of 15 genes and a locus of LPS with 3 genes, both broadly recognized as important virulence factors in HS. We find a complete phage sequence of 37,778 pb and a genomic island of 47,080 pb containing genes coding hypothetical protein.

The SC6 avian lineage contains the largest number of genes differentially present with two intact and one incomplete phage and a genomic island flanked by tRNA, the genes in this GI with unknown function. Additionally, this lineage presents a *gatYZABCR* operon responsible for galactitol metabolism in many enterobacterial and important for in vivo colonization and pathogenesis of *Salmonella* Typhimurium (Nolle et al., 2017) (Fig 5C). Next to this operon, a cluster of glucosyltransferase genes are present, the SC6 lineage doesn’t contain any known LPS genotype (Supplementary Fig. 3), so, this operon may be responsible for new LPS genotype form present only in SC6 lineage. Interestingly, a cluster of genes related to galactose metabolism including a regulatory gene and 3 other genes with unknown functions are present in the middle of *comM* gene, this insertion was mediated by transposons (Fig 5C).

The SC8 lineage associated with avian isolates has an intact phage of around 60 kb and the citrate operon *citCDEFG* (Fig 6). Citrate is ubiquitous and some microorganisms are able to metabolize it as an energy and carbon source. The structural genes *citDEF* and accessory *citC* and *citG* conform the Citrate Lyase complex to produce oxaloacetate from citrate under anaerobic conditions in *Klebsiella pneumoniae*, *Salmonella* Typhimurium and *Haemophilus influenzae* (Bott and Dimroth, 1994; Sobczak and Lolkema, 2005). Whereas SC9 lineage harbors a CRISPR cluster and a *tatD* gene flanked of hypothetical proteins. Additionally, three other operons related to carbohydrate metabolisms are present in avian lineages: L-fucose operon *fucPAUKIR*, D-allose operon *alsRBACE* and L-arabinose operon *araABCHG* (Fig 6).

**Figure 6.**
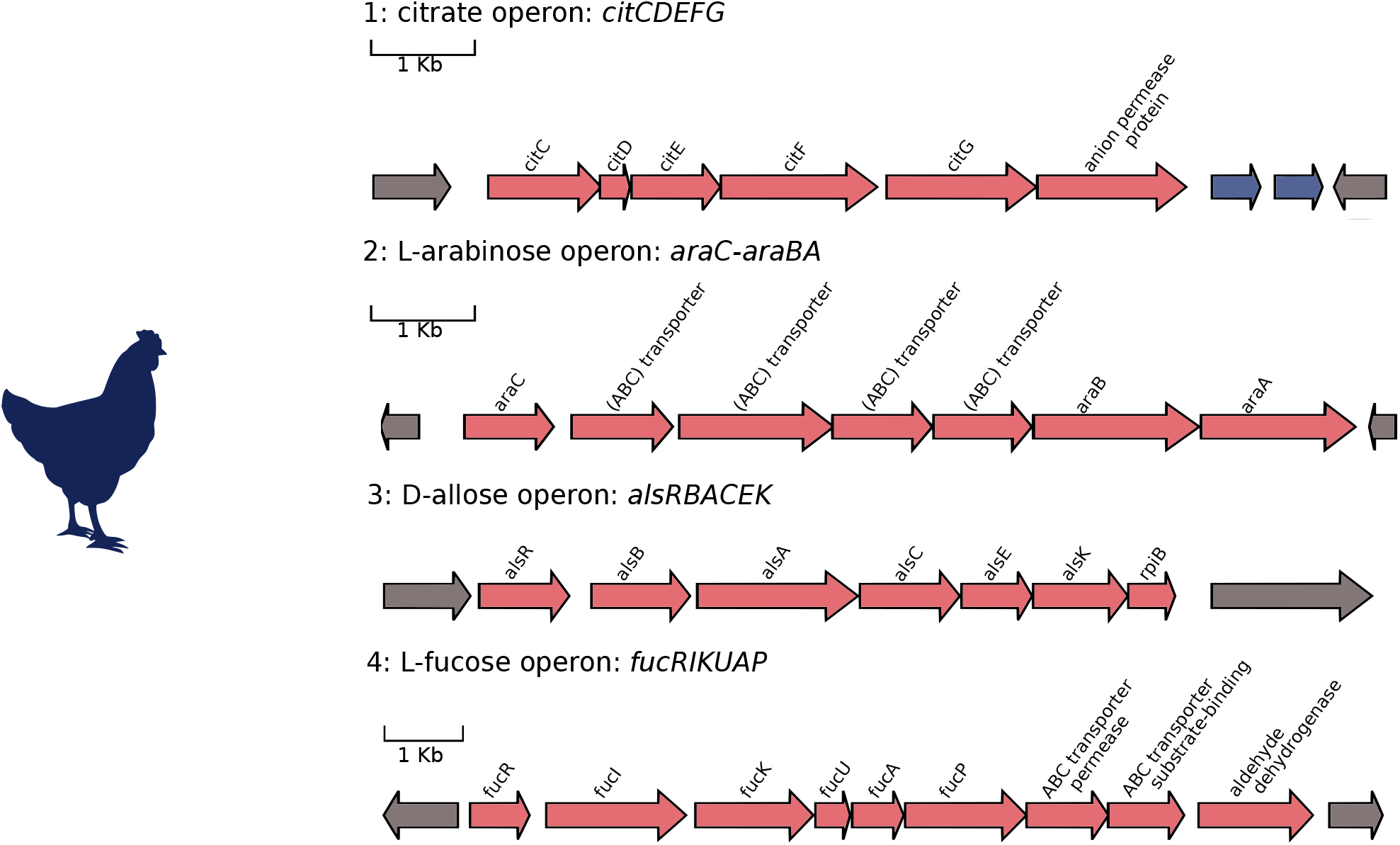
Gene annotation of carbohydrate operons related to poultry-fowl cholera isolates. Four operons related to metabolism and transport of carbohydrates are differentially presents in poultry-fowl cholera isolates. Citrate operon (citCDEFG), L-arabinose operon (*araCBA*), D-allose operon (*alsRBACEK*) and L-fucose operon (*fucRIKUAP*). These exclusive metabolic arsenal gives new metabolic capacities to facing colonization in avian hosts.

Importantly, differential gene content in distinct lineages is associated with carbohydrate transport and metabolism. This provides new energetic capacities giving a better chance to compete during pathogenesis and host colonization.

### The majority of antimicrobial resistance genes (ARGs) are present in bovine isolates

We also analyze the presence of known virulence genes and ARGs in *P. multocida* genomes, we don’t find a clear correlation between the presence of a determinate virulent gene or ARG according to host or lineage. However, some points are interesting: the complete *toxA* sequence is present in alpaca isolates and some swine strains (Fig 7). The *toxA* gene, a unique exotoxin for the species, is found in cases of progressive atrophic rhinitis in pigs belonging to serogroup D and in cases of pneumonia in different species of serogroup A (Wilson and Ho, 2013). Likewise, the *tbpA* gene was uniquely present in bovine and alpaca hosts. This gene encodes a transferrin binding protein first detected in ruminant strains (Ewers et al., 2006). Bovine strains show an increased presence of ARGs compared to strains from other hosts, mainly due to the use of antibiotics by the livestock industry increasing selection pressure to antibiotic resistance (Fig 7).

**Figure 7.**
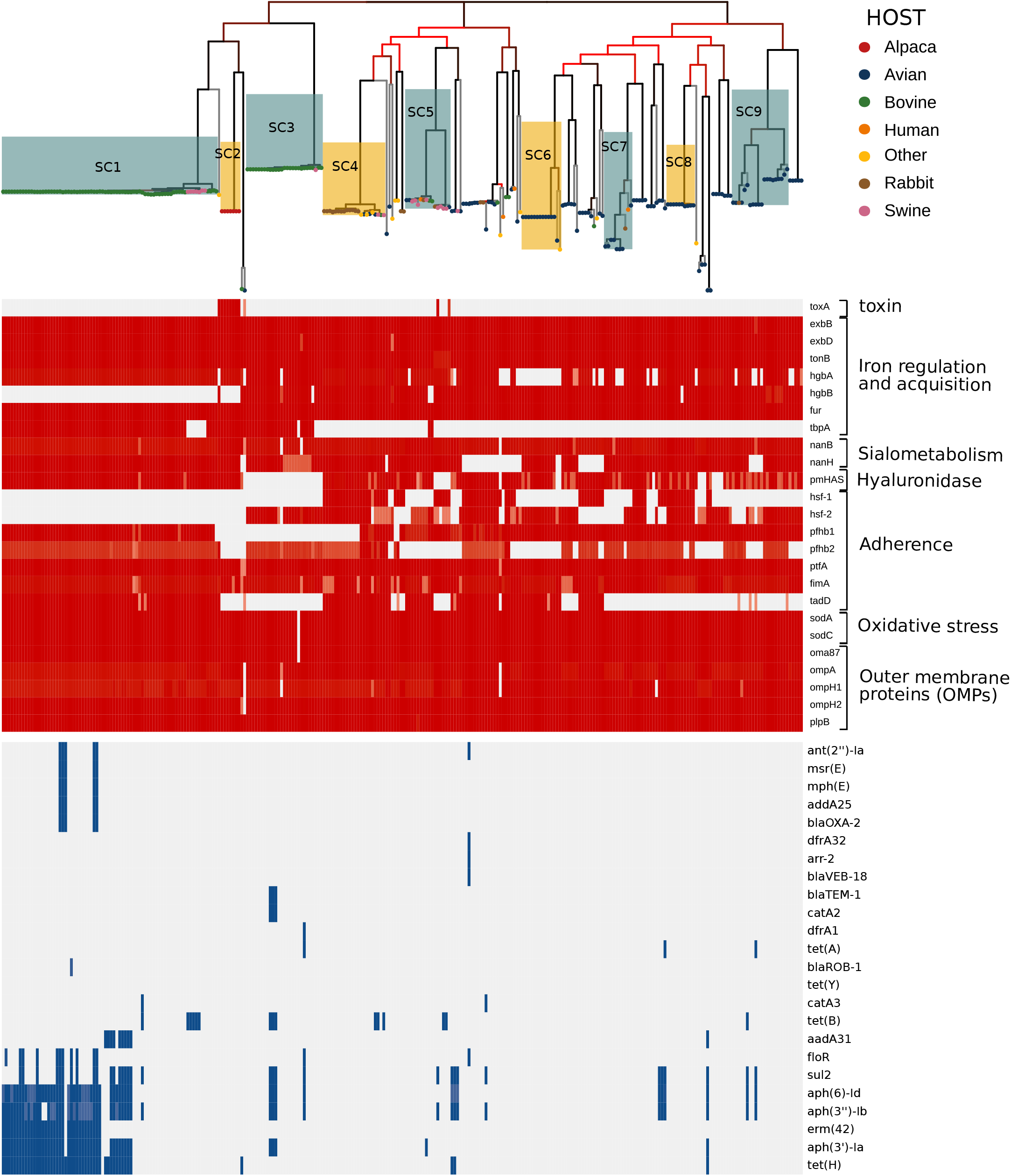
Virulence genes and antibiotic resistance genes (ARGs) in PmI *Pasteurella multocida* genome collection. ML tree of PmI phylogroups with tip-point colored according host origin and SC1-SC9 lineages are highlighted with heatmap presence/absence for virulence genes (red) and ARGs (blue).

## Discussion

The capacity to infect multiple hosts is characteristic of many bacteria and this ability is due to phenotypic plasticity and high variation in genetic content that provides ability to exploit diverse environments. *Pasteurella multocida* can infect a wide range of domestic and wild animals including humans, causing differential disease manifestation in each of them. Recent genomic studies have revealed a high diversity of genetic content in *P. multocida*, with a wide and diverse presence of plasmids, phages and genomic islands in many isolates (Peng et al., 2018; Zhu et al., 2019). Other comparative genomics studies have suggested the presence of different host-associated lineages or specific diseases with the possible presence of genetic elements unique to specific hosts (Johnson et al., 2013; Hurtado et al., 2018). Here we demonstrate the high genomic diversity and population structure of *P. multocida* from analysis of 336 complete genomes with lineages supported by phylogeny and combinations of genetic content associated with host and disease.

We report for the first time that *P. multocida* is divided into 2 clearly divergent phylogroups (PmI and PmII). The high degree of divergence and almost null inter-phylogroup recombination at the core-genome level suggests that PmI and pmII are separately evolving subpopulations and considering the nucleotide divergence (96%), can be considered as separate or evolving species (Richter and Rosselló-Móra, 2009; Jain et al., 2018). There is no experimental evidence of phenotypic difference, host restriction (isolates from both phylogroups can infect birds in the same country) or different pathogenic capacity, but the existence of barriers to genetic exchange between both phylogroups is possible. Although the presence of mechanical barriers does not seem obvious, it is likely that both populations occupy different ecological niches (cryptic) within the same host, as was previously proposed in the pathogenic bacteria *Campylobacter jejuni* (Sheppard et al., 2014). Importantly, our data show that the *torCAD* operon found in PmI is absent in PmII. The *torCAD* operon codes for a system that allows the use of TMAO as an electron acceptor during anaerobic respiration in facultative bacteria (Méjean et al., 1994). In experimental works, it has been shown that the stability of the breathing system in ETEC (Enterotoxigenic *E. coli*) is necessary for the expression of label-heat toxin (LT) under anaerobic conditions and when *torCAD* is deleted, LT is not released (Lu et al., 2016). The absence of *torCAD* has physiological effects on adaptation to anaerobic environments during colonization or invasion, and generates a different niche structure between those who have and do not have *torCAD*. This can induce physical separation between nascent lineages and, thus, a barrier to gene flow. Genotypes that carry an adaptation can initiate a speciation process by being more suitable for one habitat but less suitable for another (Wiedenbeck and Cohan, 2011).

While PmII contains isolates from birds and humans, PmI contains isolates with a larger host range. Our phylogenetic analysis of the PmI phylogroup shows high lineage diversification as previously described (Hurtado et al., 2018; Zhu et al., 2019). Additionally, analysis of population structure revealed 9 monophyletic sequence clusters (SC1-SC9) within PmI, seven of which show host and disease predilection. Some of these lineages are also concordant with Capsular: LPS: MLST genotypes as has been previously described (Peng et al., 2018). Interestingly, the accessory genome of *P. multocida* isolates from within the same SC lineage are more closely related showing that accessory gene content is heavily influenced by lineage. This highlights the importance of horizontal gene transfer in the adaptation of *P. multocida* to its host. The genetic coherence between the core and accessory genome is observed in other opportunistic pathogens such as *Staphylococcus aureus* (Richardson et al., 2018) and *E. coli* (Mcnally et al., 2016) suggesting the importance of recombination as an important force to maintain genomic cohesion in a bacterial population (Marttinen et al., 2015).

Identifying combinations of genes associated with host lineages and specific diseases reveals the wide presence of phages and genes responsible for metabolism that are most important in differentiating the gene pool. In the SC1 lineage associated with pneumonic pasteurellosis, mainly in bovines, the presence of the trehalose operon is important, while in isolates associated with poultry lineages (SC6, SC7, SC8, SC9), various carbohydrate metabolism operons such as L-fucose, D-allose, L-Arabinose, and Citrate are present conferring new metabolic capacities. In 2015, Selliey et al. reported trehalose fermentation capacity in *P. multocida* isolated from lungs of bovines with pneumonic processes but not in isolates from the nasal cavity. This suggests that additional biochemical features like trehalose metabolism could be important during colonization of the lower respiratory tract during pneumonia in bovines (Sellyei et al., 2015). Additionally, a recent study showed that the capacity of metabolize low concentrations of trehalose increase the virulence of an emergent *Clostridium difficile* ribotype in natural human infections and in a murine model (Collins et al., 2018).

The classic virulence factors identified in *P. multocida* include capsule, lipopolysaccharides, surface adhesion proteins, iron acquisition, and regulation proteins (Harper et al., 2006). However, pathogenic bacteria face a variety of environments during the infection process to which they adapt by using alternative sources of nutrients, a phenomenon called nutritional virulence (Kwaik and Bumann, 2013). *P. multocida* produces different diseases and faces different environments in each host: either intracellularly or extracellularly; in the mucosa, blood, respiratory tract, digestive tract or in the skin. The ability to obtain nutrients is a main battleground of the pathogen-host interaction and is one of the fundamental aspects of infectious diseases where pathogens face complex nutritional dynamics in host microenvironments (Rohmer et al., 2011). Expanding metabolic capacities could be a transcendental strategy to niche adaptation considering that glucose concentration outside cells is low with many other carbohydrates widely abundant (Passalacqua et al., 2016). In avian cholera, the primary entry for *P. multocida* occurs through the airways to the lungs. However, the survival of different strains of *P. multocida* in the intestines, and subsequent colonization of the ileum and jejunum mucosa, gives rise to the possibility of entry through the digestive tract as an alternative route (Lee et al., 2000; Mbuthia et al., 2011). These strains act at an extracellular level in the respiratory and intestinal mucosa, in which the energy requirements are variable, even facing the resident microbiota. Therefore, the additional presence of genes and operons with carbohydrate metabolism and transport functions in the accessory genome can promote the colonization and establishment of strains of *P. multocida* associated with avian cholera on the surface of respiratory or intestinal tissue.

Together, our data provide detailed genomic information about population structure and accessory genome diversity in *P. multocida*. We present strong evidence of two highly divergent nonrecombinat phylogroups in *P. multocida* (Pml and Pmll), potentially driven by the *torCAD* operon which is critically involved in anaerobic respiration and present only in PmI phylogroup. Furthermore, we found accessory gene pools correlated to lineages associated to host and disease predilection. Interestingly, some of these gene pools are carbohydrate metabolism operons present differentially in bovine and avian lineages. These findings suggest that additional metabolic pathways could be important during colonization conducting a host adaptation of these lineages. Further experimental studies are necessary in order to confirm differential metabolic capacities among strains from different lineages.

## Acknowledgements

This study was supported by the Programa Nacional de Innovación para la Competitividad y Productividad (Innovate Peru), contracts 133-FINCyT-IB-2013 and 288-INNOVATEPERU-EC-2017; and the Programa de Promoción de Tesis de Pregrado 2017 (Project Number: A17080054b) from the Vicerrectorado de Investigación y Postgrado, Universidad Nacional Mayor de San Marcos (VRIP-UNMSM).

## Author Contribution

DC conceived the idea and designed the experiments. RR, LL and LM collected and provided samples. DC analyzed the data. DC and RH wrote the manuscript. DC, RH, LL, JCW, RR, VA and LM Review & Editing, VA and LM supervised this work. All authors revised and approved the manuscript previous to submission.

## Supplementary figure legends

**Supplementary fig 1.**
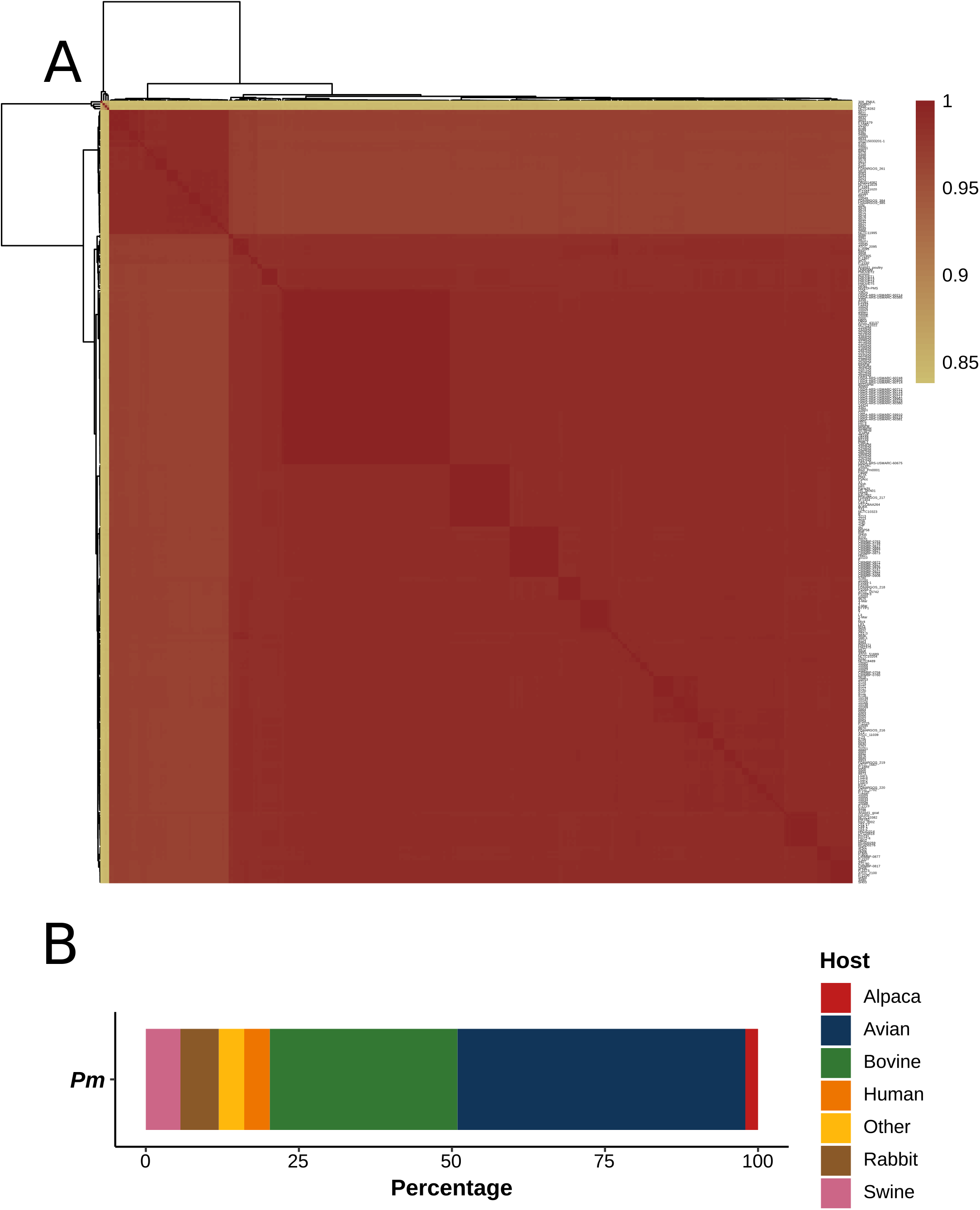
Specie membership of 340 genomes putative *P. multocida* and final count of 336 *P. multocida* isolates (>95% identity). (A) Hierarchical clustering for 340 genomes according ANI values. 4 genomes (<85% ANI) were removes for subsequent analysis. (B) Barplot of host distribution for 336 *P. multocida* genomes (> 95% ANI).

**Supplementary fig 2.**
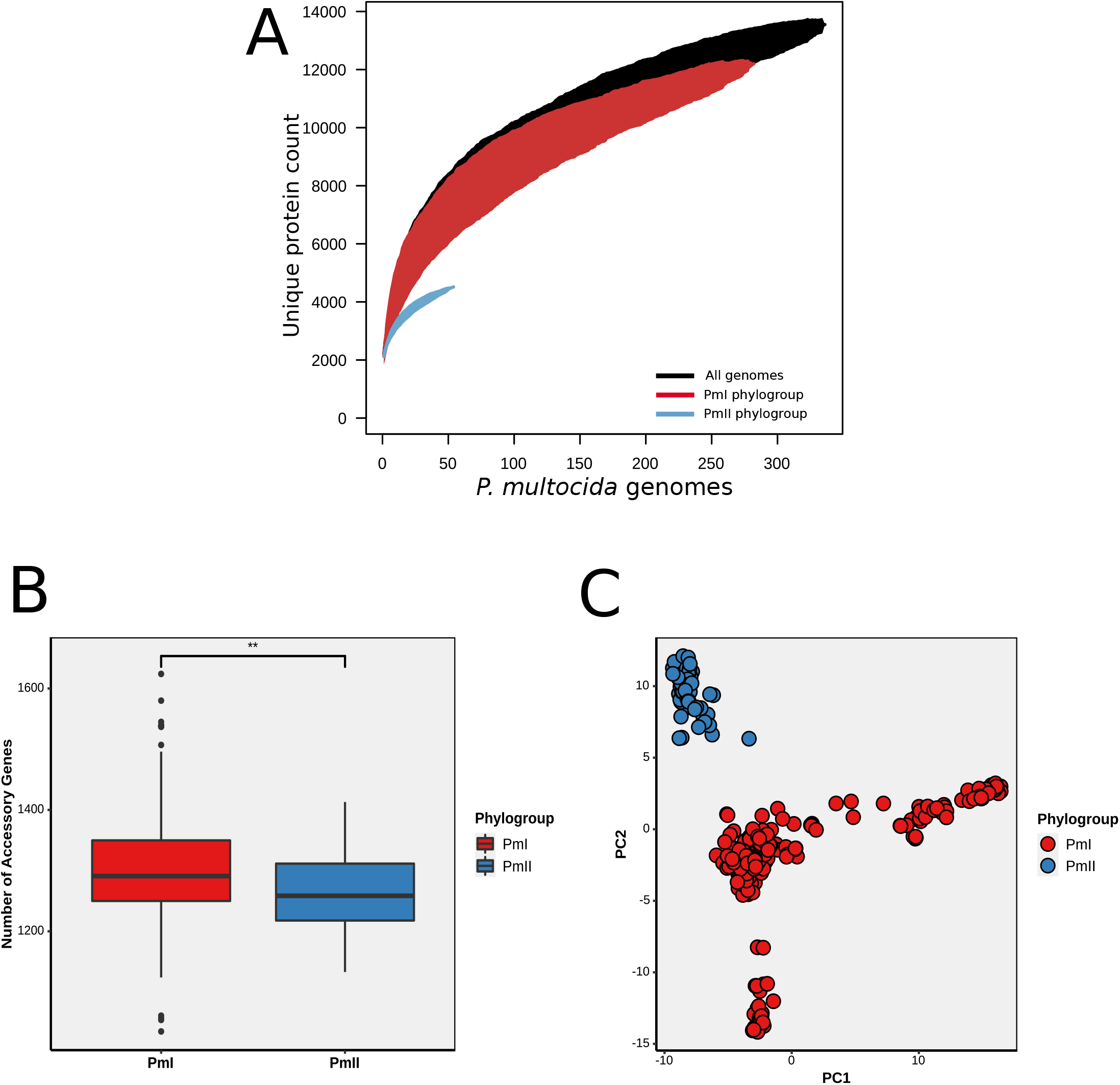
Pangenome and accessory genome distribution in PmI and PmII phylogroups. (A) Pangenome accumulation curves for *P. multocida* full collection (black) and for PmI (red) and PmII (blue) phylogroups. (B) Boxplots showing the number of accessory genes in both PmI and PmII phylogroups. PmI possesses a slightly significantly bigger accessory genome than PmII phylogroup. (p = 0.00168, Mann Whitney U test). (C) PCA analysis based on the presence of common (1–99% prevalence) accessory genes.

**Supplementary fig 3.**
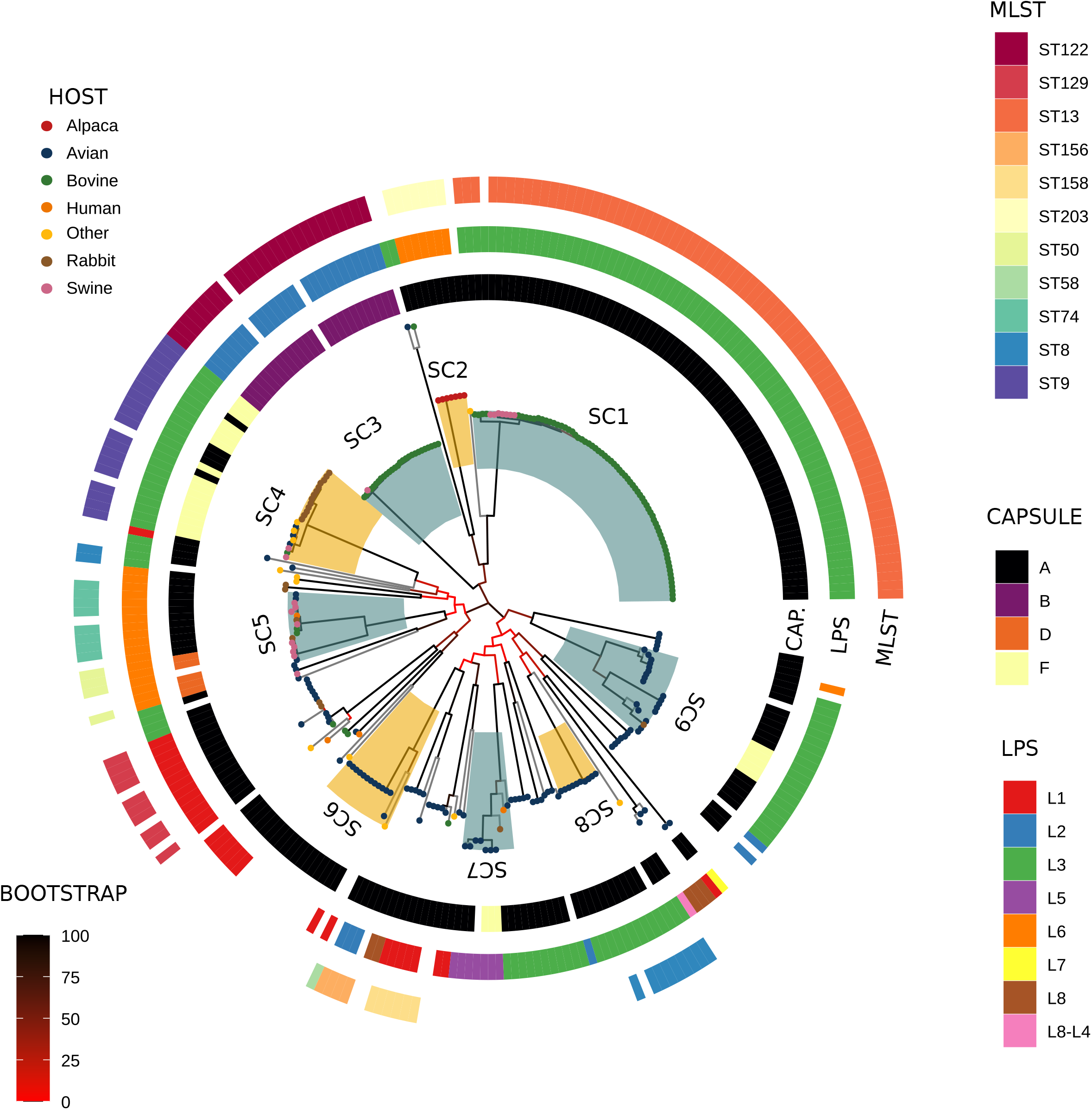
Maximum likelihood tree of 282 isolates from PmI phylogroup with LPS, capsular and MLST annotation. ML tree and population structure of PmI phylogroups shows nine monophyletic sequence clusters (SCs). LPS and Capsular genotypes annotation for each isolate are showed in rings around to the phylogeny. Branch colors indicate bootstrap support according to the legend provided in the figure. hierBAPS sequence clusters are highlighted in alternating colors and labeled.

**Supplementary fig 4.**
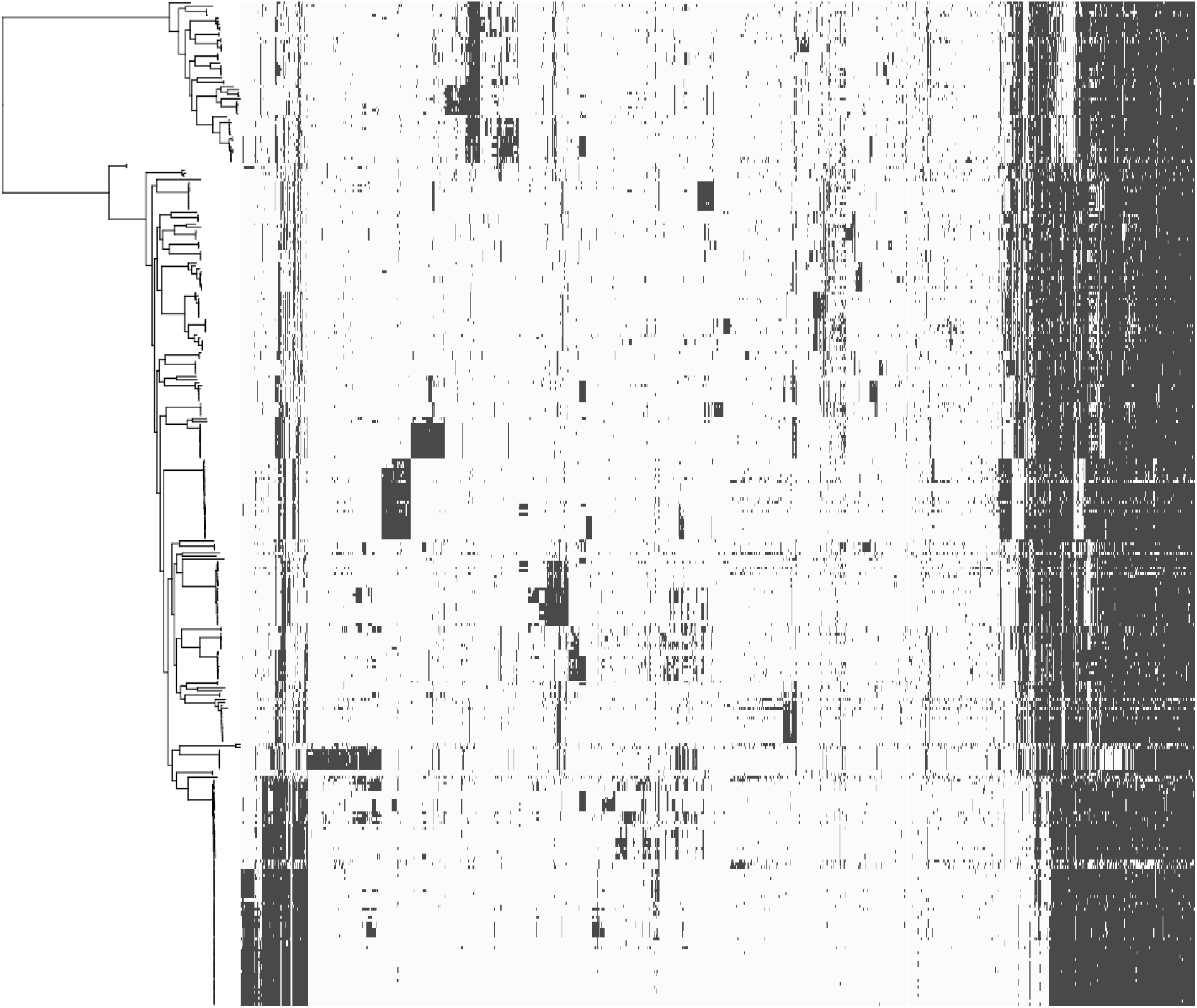
Accessory gene distribution in 366 *P. multocida* genomes. Heatmap of 5152 accessory genes presents (black) or absents (white) in 336 *P. multocida* genomes shows the presence of differential gene content.

**Supplementary fig 5.**
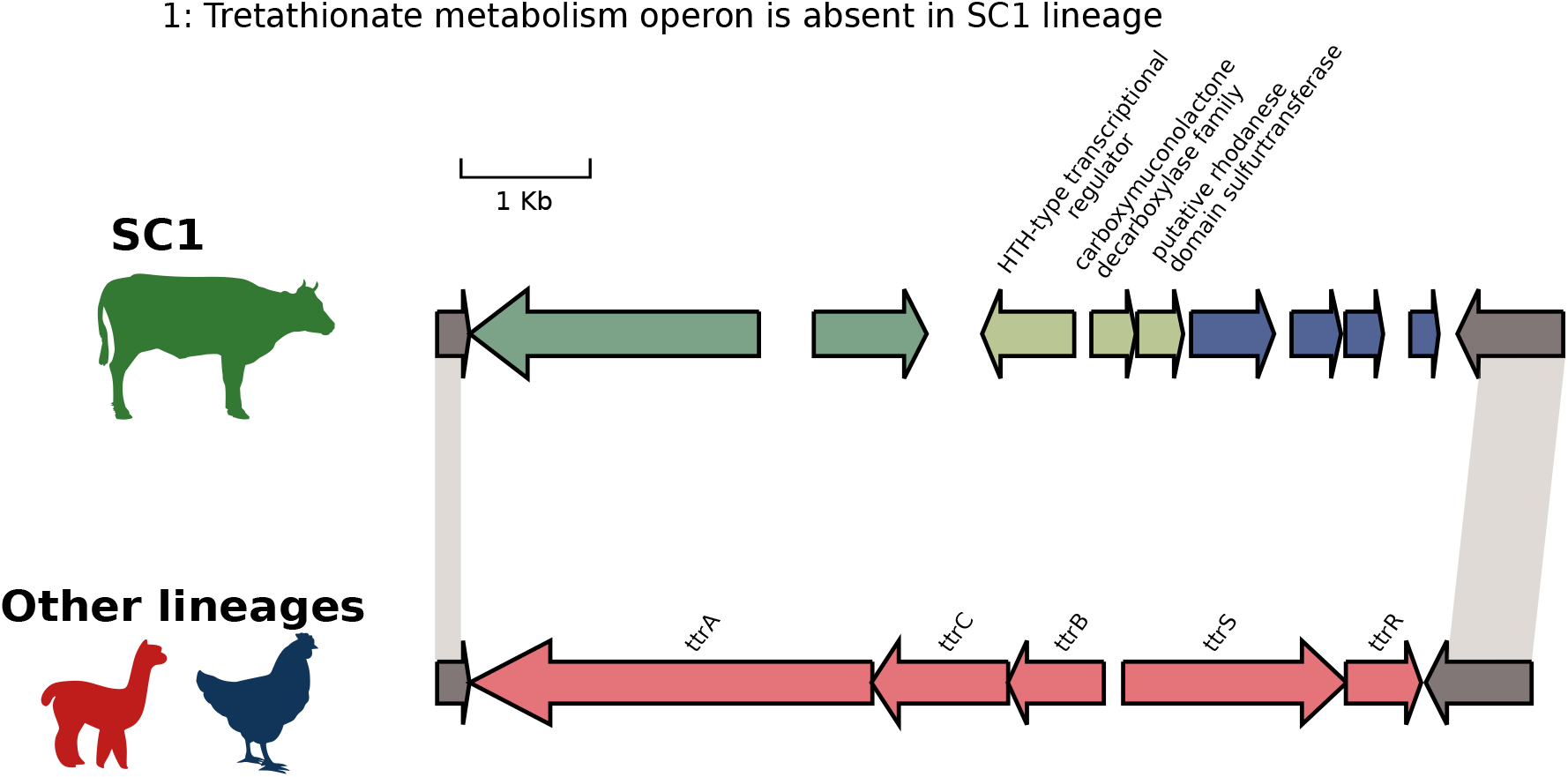
Gene annotation of gene cluster absent in SC1 lineage associate with pneumonia in bovines. Gene annotation for a Tretathionate metabolism operon present in all lineages but absent in SC1 lineage.

## References

Alcock, B. P., Raphenya, A. R., Lau, T. T. Y., Tsang, K. K., Bouchard, M., Edalatmand, A., et al. (2020). CARD 2020: Antibiotic resistome surveillance with the comprehensive antibiotic resistance database. Nucleic Acids Res. 48, D517–D525. doi:10.1093/nar/gkz935.

Bankevich, A., Nurk, S., Antipov, D., Gurevich, A. A., Dvorkin, M., Kulikov, A. S., et al. (2012). SPAdes: A new genome assembly algorithm and its applications to single-cell sequencing. J. Comput. Biol. 19, 455–477. doi:10.1089/cmb.2012.0021.

Bolger, A. M., Lohse, M., and Usadel, B. (2014). Trimmomatic: A flexible trimmer for Illumina sequence data. Bioinformatics 30, 2114–2120. doi:10.1093/bioinformatics/btu170.

Bott, M., and Dimroth, P. (1994). Klebsiella pneumoniae genes for citrate lyase and citrate lyase ligase: Localization, sequencing, and expression. Mol. Microbiol. 14, 347–356. doi:10.1111/j.1365-2958.1994.tb01295.x.

Boyce, J. D., and Adler, B. (2000). The Capsule Is a Virulence Determinant in the Pathogenesis of Pasteurella multocida M1404 (B:2). Infect. Immun. 68, 3463–3468.

Boyce, J. D., Seemann, T., Adler, B., and Harper, M. (2012). Pathogenomics of Pasteurella multocida. Curr. Top. Microbiol. Immunol. 361, 23–38. doi:10.1007/82-2012-203.

Chung, J. Y., Wilkie, I., Boyce, J. D., and Adler, B. (2001). Vaccination against fowl cholera with acapsular pasteurella multocida A:1. Infect. Immun. 69, 2487–2492. doi:10.1128/IAI.69.4.2487-2492.2001.

Collins, J., Robinson, C., Danhof, H., Knetsch, C. W., van Leeuwen, H. C., Lawley, T. D., et al. (2018). Dietary trehalose enhances virulence of epidemic Clostridium difficile. Nature 553, 291–294. doi:10.1038/nature25178.

Croucher, N. J., Page, A. J., Connor, T. R., Delaney, A. J., Keane, J. A., Bentley, S. D., et al. (2015). Rapid phylogenetic analysis of large samples of recombinant bacterial whole genome sequences using Gubbins. Nucleic Acids Res. 43, e15. doi:10.1093/nar/gku1196.

Díaz, E., Ferrández, A., Prieto, M. A., and García, J. L. (2001). Biodegradation of Aromatic Compounds byEscherichia coli. Microbiol. Mol. Biol. Rev. 65, 523–569. doi:10.1128/mmbr.65.4.523-569.2001.

Ewers, C., Lübke-Becker, A., Bethe, A., Kießling, S., Filter, M., and Wieler, L. H. (2006). Virulence genotype of Pasteurella multocida strains isolated from different hosts with various disease status. Vet. Microbiol. 114, 304–317. doi:10.1016/j.vetmic.2005.12.012.

Fàbrega, A., and Vila, J. (2013). Salmonella enterica serovar Typhimurium skills to succeed in the host: Virulence and regulation. Clin. Microbiol. Rev. 26, 308–341. doi:10.1128/CMR.00066-12.

Gupta, S. K., Padmanabhan, B. R., Diene, S. M., Lopez-Rojas, R., Kempf, M., Landraud, L., et al. (2014). ARG-annot, a new bioinformatic tool to discover antibiotic resistance genes in bacterial genomes. Antimicrob. Agents Chemother. 58, 212–220. doi:10.1128/AAC.01310-13.

Harper, M., Boyce, J. D., and Adler, B. (2006). Pasteurella multocida pathogenesis: 125 Years after Pasteur. FEMS Microbiol. Lett. 265, 1–10. doi:10.1111/j.1574-6968.2006.00442.x.

Harper, M., Boyce, J. D., and Adler, B. (2012). “The key surface components of Pasteurella multocida: Capsule and lipopolysaccharide,” in Pasteurella multocida, 39–51. doi:10.1007/82-2012-202.

Harper, M., Cox, A. D., Michael, F. S., Wilkie, I. W., Boyce, J. D., and Adler, B. (2004). A Heptosyltransferase Mutant of Pasteurella multocida Produces a Truncated Lipopolysaccharide Structure and Is Attenuated in Virulence. Infect. Immun. 72, 3436–3443.

Harper, M., John, M., Turni, C., Edmunds, M., St. Michael, F., Adler, B., et al. (2015). Development of a rapid multiplex PCR assay to genotype pasteurella multocida strains by use of the lipopolysaccharide outer core biosynthesis locus. J. Clin. Microbiol. 53, 477–485. doi:10.1128/JCM.02824-14.

Harrison, K. J., Crécy-Lagard, V. De, and Zallot, R. (2018). Gene Graphics: A genomic neighborhood data visualization web application. Bioinformatics 34, 1406–1408. doi:10.1093/bioinformatics/btx793.

Hotchkiss, E. J., Hodgson, J. C., Lainson, A., and Zadoks, R. N. (2011a). Multilocus sequence typing of a global collection of Pasteurella multocida isolates from cattle and other host species demonstrates niche association. BMC Microbiol. 11, 1–8. doi:10.1186/1471-2180-11-115.

Hotchkiss, E. J., Hodgson, J. C., Leemput, E. S. De, Dagleish, M. P., and Zadoks, R. N. (2011b). Molecular epidemiology of Pasteurella multocida in dairy and beef calves. Vet. Microbiol. 151, 329–335. doi:10.1016/j.vetmic.2011.03.018.

Huerta-Cepas, J., Forslund, K., Coelho, L. P., Szklarczyk, D., Jensen, L. J., Von Mering, C., et al. (2017). Fast genome-wide functional annotation through orthology assignment by eggNOG-mapper. Mol. Biol. Evol. 34, 2115–2122. doi:10.1093/molbev/msx148.

Hurtado, R., Carhuaricra, D., Soares, S., Viana, M. V. C., Azevedo, V., Maturrano, L., et al. (2018). Pan-genomic approach shows insight of genetic divergence and pathogenic-adaptation of Pasteurella multocida. Gene 670, 193–206. doi:10.1016/j.gene.2018.05.084.

Hurtado, R., Maturrano, L., Azevedo, V., and Aburjaile, F. (2020). Pathogenomics insights for understanding Pasteurella multocida adaptation. Int. J. Med. Microbiol. 310, 151417. doi:10.1016/j.ijmm.2020.151417.

Jain, C., Rodriguez-R, L. M., Phillippy, A. M., Konstantinidis, K. T., and Aluru, S. (2018). High throughput ANI analysis of 90K prokaryotic genomes reveals clear species boundaries. Nat. Commun. 9, 1–8. doi:10.1038/s41467-018-07641-9.

Johnson, T. J., Abrahante, J. E., Hunter, S. S., Hauglund, M., Tatum, F. M., Maheswaran, S. K., et al. (2013). Comparative genome analysis of an avirulent and two virulent strains of avian Pasteurella multocida reveals candidate genes involved in fitness and pathogenicity. BMC Microbiol. 13, 1–10. doi:10.1186/1471-2180-13-106.

Kloepper, T. H., and Huson, D. H. (2008). Drawing explicit phylogenetic networks and their integration into SplitsTree. BMC Evol. Biol. 8, 22. doi:10.1186/1471-2148-8-22.

Kwaik, Y. A., and Bumann, D. (2013). Microreview Microbial quest for food in vivo: ‘Nutritional virulence’ as an emerging paradigm. Cell. Microbiol. 15, 882–890. doi:10.1111/cmi.12138.

Lee, C. W., Wilkie, I. W., Townsend, K. M., and Frost, A. J. (2000). The demonstration of Pasteurella multocida in the alimentary tract of chickens after experimental oral infection. Vet. Microbiol. 72, 47–55.

Liu, C., Zheng, H., Yang, M., Xu, Z., Wang, X., Wei, L., et al. (2015). Genome analysis and in vivo virulence of porcine extraintestinal pathogenic Escherichia coli strain PCN033. BMC Genomics 16, 717. doi:10.1186/s12864-015-1890-9.

Lu, X., Fu, E., Xie, Y., and Jin, F. (2016). Electron acceptors induce secretion of enterotoxigenic Escherichia coli heat-labile enterotoxin under anaerobic conditions through promotion of GspD assembly. Infect. Immun. 84, 2748–2757. doi:10.1128/IAI.00358-16.

Marttinen, P., Croucher, N. J., Gutmann, M. U., Corander, J., and Hanage, W. P. (2015). Recombination produces coherent bacterial species clusters in both core and accessory genomes. Microb. Genomics 1, e000038. doi:10.1099/mgen.0.000038.

Mbuthia, P. G., Njagi, L. W., Nyaga, P. N., Bebora, L. C., Minga, U., Christensen, J. P., et al. (2011). Time-course investigation of infection with a low virulent Pasteurella multocida strain in normal and immune-suppressed 12-week-old free-range chick. Avian Pathol. 40, 629–637. doi:10.1080/03079457.2011.623298.

Mcnally, A., Oren, Y., Kelly, D., Pascoe, B., Dunn, S., Sreecharan, T., et al. (2016). Combined Analysis of Variation in Core, Accessory and Regulatory Genome Regions Provides a Super-Resolution View into the Evolution of Bacterial Populations. PLoS Genet. 12, e1006280. doi:10.5061/dryad.d7d71.

Méjean, V., lobbi-Nivol, C., Lepelletier, M., Giordano, G., Chippaux, M., and Pascal, M.-C. (1994). TMAO anaerobic respiration in Escherichia coli: involvement of the tor operon. Mol. Microbiol. 11, 1169–1179. doi:10.1111/j.1365-2958.1994.tb00393.x.

Michael, G. B., Kadlec, K., Sweeney, M. T., Brzuszkiewicz, E., Liesegang, H., Daniel, R., et al. (2012). ICEPmu1, an integrative conjugative element (ICE) of Pasteurella multocida: structure and transfer. J. Antimicrob. Chemother. 67, 91–100. doi:10.1093/jac/dkr411.

Moustafa, A. M., Seemann, T., Gladman, S., Adler, B., Harper, M., Boyce, J. D., et al. (2015). Comparative genomic analysis of Asian haemorrhagic septicaemia-associated strains of Pasteurella multocida identifies more than 90 haemorrhagic septicaemia-specific genes. PLoS One 10. doi:10.1371/journal.pone.0130296.

Nolle, N., Felsi, A., Heermann, R., and Fuchs, T. M. (2017). Genetic Characterization of the Galactitol Utilization Pathway of Salmonella enterica Serovar Typhimurium. J. Bacteriol. 199, 1–16.

Page, A. J., Cummins, C. A., Hunt, M., Wong, V. K., Reuter, S., Holden, M. T. G., et al. (2015). Roary: Rapid large-scale prokaryote pan genome analysis. Bioinformatics 31, 3691–3693. doi:10.1093/bioinformatics/btv421.

Passalacqua, K. D., Charbonneau, M., and Riordan, M. X. D. O. (2016). Bacterial Metabolism Shapes the Host – Pathogen Interface. Microbiol. Spectr. 4, 1–21. doi:10.1128/microbiolspec.VMBF-0027-2015.Correspondence.

Peng, Z., Liang, W., Wang, F., Xu, Z., Xie, Z., Lian, Z., et al. (2018). Genetic and Phylogenetic Characteristics of Pasteurella multocida Isolates From Different Host Species. Front. Microbiol. 9, 1408. doi:10.3389/fmicb.2018.01408.

Peng, Z., Wang, X., Zhou, R., Chen, H., Wilson, B. A., and Wu, B. (2019). Pasteurella multocida: Genotypes and Genomics. Microbiol. Mol. Biol. Rev. 83. doi:10.1128/mmbr.00014-19.

Pessia, A., Grad, Y., Cobey, S., Puranen, J. S., and Corander, J. (2015). K-Pax2: Bayesian identification of cluster-defining amino acid positions in large sequence datasets. Microb. Genomics 1, e000025. doi:10.1099/mgen.0.000025.

Petersen, A., Bisgaard, M., Townsend, K., and Christensen, H. (2014). MLST typing of Pasteurella multocida associated with haemorrhagic septicaemia and development of a real-time PCR specific for haemorrhagic septicaemia associated isolates. Vet. Microbiol. 170, 335–341. doi:10.1016/j.vetmic.2014.02.022.

Richardson, E. J., Bacigalupe, R., Harrison, E. M., Weinert, L. A., Lycett, S., Vrieling, M., et al. (2018). Gene exchange drives the ecological success of a multi-host bacterial pathogen. Nat. Ecol. Evol. 2, 1468–1478. doi:10.1038/s41559-018-0617-0.

Richter, M., and Rosselló-Móra, R. (2009). Shifting the genomic gold standard for the prokaryotic species definition. Proc. Natl. Acad. Sci. U. S. A. 106, 19126–19131. doi:10.1073/pnas.0906412106.

Rímac, R., Luna, L., Hurtado, R., Rosadio, R., and Maturrano, L. (2017). Detection and genetic characterization of Pasteurella multocida from alpaca (Vicugna pacos) pneumonia cases. Trop. Anim. Heal. Prod. 49, 1325–1328. doi:10.1007/s11250-017-1309-5.

Rohmer, L., Hocquet, D., and Miller, S. I. (2011). Are pathogenic bacteria just looking for food? Metabolism and microbial pathogenesis. Trends Microbiol. 19, 341–348. doi:10.1016/j.tim.2011.04.003.

Rosadio, R., Cirilo, E., Manchego, A., and Rivera, H. (2011). Respiratory syncytial and parainfluenza type 3 viruses coexisting with Pasteurella multocida and Mannheimia hemolytica in acute pneumonias of neonatal alpacas. Small Rumin. Res. 97, 110–116. doi:10.1016/j.smallrumres.2011.02.001.

Seemann, T. (2014). Prokka: rapid prokaryotic genome annotation. Bioinformatics 30, 2068–2069. doi:10.1093/bioinformatics/btu153.

Sellyei, B., Rónai, Z. S., Jánosi, S., and Makrai, L. (2015). Comparative analysis of pasteurella multocida strains isolated from bovine respiratory infections. Acta Microbiol. Immunol. Hung. 62, 453–461. doi:10.1556/030.62.2015.4.9.

Sheppard, S. K., Cheng, L., Méric, G., de Haan, C. P. A., Llarena, A.-K., Marttinen, P., et al. (2014). Cryptic ecology among host generalist *Campylobacter jejuni* in domestic animals. Mol. Ecol. 23, 2442–2451. doi:10.1111/mec.12742.

Shivachandra, S. B., Viswas, K. N., and Kumar, A. A. (2011). A review of hemorrhagic septicemia in cattle and buffalo. Anim. Heal. Res. Rev. 12, 67–82. doi:10.1017/S146625231100003X.

Sobczak, I., and Lolkema, J. S. (2005). The 2-Hydroxycarboxylate Transporter Family: Physiology, Structure, and Mechanism. Microbiol. Mol. Biol. Rev. 69, 665–695. doi:10.1128/MMBR.69.4.665.

Stamatakis, A. (2014). RAxML version 8: a tool for phylogenetic analysis and post-analysis of large phylogenies. Bioinformatics 30, 1312–1313. doi:10.1093/bioinformatics/btu033.

Tonkin-Hill, G., Lees, J. A., Bentley, S. D., Frost, S. D. W., and Corander, J. (2018). RhierBAPs: An R implementation of the population clustering algorithm hierbaps [version 1; referees: 2 approved]. Wellcome Open Res. 3, 93. doi:10.12688/wellcomeopenres.14694.1.

Townsend, K. M., Boyce, J. D., Chung, J. Y., Frost, A. J., and Adler, B. (2001). Genetic organization of Pasteurella multocida cap loci and development of a multiplex capsular PCR typing system. J. Clin. Microbiol. 39, 924–929. doi:10.1128/JCM.39.3.924-929.2001.

Wiedenbeck, J., and Cohan, F. M. (2011). transfer and adaptation to new ecological niches. FEMS Microbiol. Lett. 35, 957–976. doi:10.1111/j.1574-6976.2011.00292.x.

Wilkie, I. W., Harper, M., Boyce, J. D., and Adler, B. (2012). Pasteurella multocida: Diseases and pathogenesis. Curr. Top. Microbiol. Immunol. 361, 1–22. doi:10.1007/82-2012-216.

Wilson, B. A., and Ho, M. (2013). Pasteurella multocida: from zoonosis to cellular microbiology. Clin. Microbiol. Rev. 26, 631–55. doi:10.1128/CMR.00024-13.

Winter, S. E., Thiennimitr, P., Winter, M. G., Butler, B. P., Huseby, D. L., Crawford, R. W., et al. (2010). Gut inflammation provides a respiratory electron acceptor for Salmonella. Nature 467, 426–429. doi:10.1038/nature09415.

Zhu, D., He, J., Yang, Z., Wang, M., Jia, R., Chen, S., et al. (2019). Comparative analysis reveals the Genomic Islands in Pasteurella multocida population genetics: On Symbiosis and adaptability. BMC Genomics 20, 63. doi:10.1186/s12864-018-5366-6.

